# Three-dimensional Genome Organization Maps in Normal Haematopoietic Stem Cells and Acute Myeloid Leukemia

**DOI:** 10.1101/2020.04.18.047738

**Authors:** Benny Wang, Lingshi Kong, Deepak Babu, Ruchi Choudhary, Winnie Fam, Jia Qi Tng, Yufen Goh, Bertrand Wong Jern Han, Xin Liu, Fang Fang Song, Priscella Chia, Ming Chun Chan, Omer An, Cheng Yong Tham, Touati Benoukraf, Henry Yang, Wilson Wang, Wee Joo Chng, Daniel Tenen, Melissa Jane Fullwood

## Abstract

Acute Myeloid Leukemia (AML) is a highly lethal blood cancer arising due to aberrant differentiation of haematopoietic stem cells. Here we obtained 3D genome organization maps by Hi-C in the CD34+ haematopoietic stem cells from three healthy individuals and eight individuals with AML, and found that AML have increased loops to oncogenes compared with normal CD34+ cells. The *MEIS1* oncogenic transcription factor is regulated by a Frequently Interacting Region (FIRE). This FIRE is only present in normal bone marrow samples, and four of eight AML sample. FIRE presence is associated with *MEIS1* expression. CRISPR excision of a FIRE boundary led to loss of *MEIS1* and reduced cell growth. Moreover, MEIS1 can bind to the promoter of *HOXA9*, and HOXA9 shows gain of Acute Myeloid Leukemia-specific super-enhancers that loop to the *HOXA9* promoter.

**Significance:** We found that Acute Myeloid Leukemias have more chromatin loops to oncogenes compared with normal blood stem cells. We identified heterogeneity in chromatin interactions at oncogenes, and heterogeneity in super-enhancers that loop to oncogenes, as two key epigenetic mechanisms that underlie *MEIS1* and *HOXA9* oncogene expression respectively.

## Introduction

Acute Myeloid Leukemia (AML) is a highly lethal cancer which is characterized by a block in the differentiation and abnormal proliferation of hematopoietic progenitor cells (1). While complete remissions in patients are generally achieved, relapses are commonly observed and eventually fatal (2,3). The DNA mutation load of AML is lower than that of solid cancers, and analyses of the mutated genes have revealed that epigenetic factors and transcription factors are frequently mutated in AML (4,5), pointing to epigenetic dysregulation as being important to AML pathogenesis. HOXA9 and MEIS1 oncogenic transcription factors are overexpressed in more than half of AML cases (6–9), and overexpression of *HOXA9* and *MEIS1* oncogenes is associated with poor prognosis (9–11). Here we asked whether there are epigenetic mechanisms behind the heterogeneity of oncogene expression in AML.

One potential source of heterogeneity is that of 3D genome organization. The genome is organized into chromatin interactions, which refer to two or more separate genome loci that come together in close spatial proximity (12), such as enhancer-promoter loops and also large domains of interacting regions, called Topologically-Associated Domains (TADs). Clusters of enhancers marked by high H3K27ac signals called super-enhancers (SEs) can be acquired by cancer cells, and have been identified to be associated with oncogene activation in cancer (13). We and others have found that SEs can loop via chromatin interactions to distant oncogenes (14). Moreover, cancer cells can show altered TADs (15) and altered chromatin loops at key oncogenes such as *TERT* (16,17). Thus, we reasoned that the interplay between SEs and chromatin interactions might underlie the heterogeneity of *HOXA9* and *MEIS1* expression in AML.

Here we performed Hi-C to obtain 3-dimensional genome organization maps in eight samples of AML bone marrow clinical samples, and three CD34+ enriched bone marrow samples from healthy individuals. The AMLs showed more loops to oncogenes than the normal blood stem cells. Interestingly, the *MEIS1* locus shows a “Frequently Interacting Region” (FIRE) that can be found in all healthy bone marrow samples, but only in four of the eight AML patient samples. FIREs are regions of unusually high local contact frequency that tend to be depleted near Topologically-Associated Domain (TAD) boundaries and enriched towards the centre of TADs (18). From the RNA-Seq of the samples, we observed that AML patient with intact *MEIS1* FIRE have *MEIS1* gene expression, but this was absent in the other samples that lacked the *MEIS1* FIRE. Removal of a FIRE boundary at the *MEIS1* gene locus leads to loss of *MEIS1* gene expression, slower leukemic cell growth and reorganisation of existing chromatin loops. *HOXA9*, in contrast to *MEIS1*, shows chromatin interactions that are present in all AML patient samples and all healthy samples, however, analysis of a large dataset of H3K27ac ChIP-Seq in AML patient samples (19) indicates the loci that interact with the *HOXA9* show differences in occupancy by SEs. Patient samples that contain SEs at the interacting loci show higher *HOXA9* expression.

Taken together, in this paper, we obtained a resource of Hi-C maps in AML and normal blood stem cells, which showed that AMLs have more loops to oncogenes than normal blood stem cells. *MEIS1* gene expression level and cell growth was dependent on maintenance of a key FIRE boundary, indicating that changes in chromatin interactions could affect gene expression and cancer cell survival. We identified two key epigenetic mechanisms underlying the heterogeneity of *MEIS1* and *HOXA9* oncogene expression in AML. First, heterogeneity of chromatin interaction structures in AML cells can explain the heterogeneous expression of *MEIS1* in AML. Second, heterogeneity of AML-acquired SEs at genomic loci that show chromatin interactions to *HOXA9* can explain heterogeneous expression of *HOXA9* in AML. This information could be useful in the development of targeted epigenetic therapies against *HOXA9* and *MEIS1* in AML while sparing healthy cell types such as normal haematopoietic stem cells.

## Results

### Hi-C analysis indicates that Acute Myeloid Leukemias have more loops to oncogenes than normal blood stem cells

To date there is no detailed study of topologically associated domains (TADs) and chromatin loops in AML as compared with normal haematopoietic stem cells, which are the putative precursor cells for AML development. Here we performed deep sequencing Hi-C in eight samples of AML bone marrow clinical samples (“AD796”, “AD903”, “AML28”, “AML29”, “AML30”, “AML42”, “AML43”and “AML44”) and three CD34+-enriched bone marrow samples from healthy individuals obtained from knee replacement surgery (“Femur 47”, “Femur 49”, “Femur 50”) (**Table S1**). AML28, 29,30, 42, 43 and 44 were obtained from fresh isolates. AD796 and AD903 were obtained from frozen stocks of AML patient samples. We note that the level of CD34+ cells in some of these patients are quite high at (**Table S1**), and therefore cannot rule out the possibility that there may be several pre-malignant changes such as clonal haematopoiesis and early myelodysplastic syndromes. All samples were obtained with patient consent.

We obtained more than a billion sequencing reads for each Hi-C library and analysed the data by Juicer (20) (**Table S2**). TADs were called using Arrowhead (20) (**Table S3**), while loops were called using HiCCUPS (20) (**Table S4**). Chromosomal heatmaps of the samples were visualized with Juicebox (**Figure S1A& B**) (21). The TADs are observed to be distinct in each of this samples. Several thousands of TADs and over ten thousand loops were called from our Hi-C data, reflecting that our libraries are of high resolution (AML28: 1,682 TADs and 24,394 loops; AML29: 1,108 TADs and 19,541 loops; AML30: 1,296 TADs and 10,733 loops; Femur47: 1,153 TADs and 4,733 loops; Femur49: 927 TADs and 13,107 loops; Femur50: 1,234 TADs and 13,795 loops).

We approached this by performing principal component analysis on three AML CD34+ (“AML28”, “AML29”, “AML30”) and three normal femur CD34+ samples (“Femur47”, “Femur 49”, “Femur 50”) to examine if they their gene expression profiles are distinctly different from each other (**Figure 1A & 2A**). We investigated the extent of similarity between loops in AML28, 29, 30 and femurs 47,49 & 50 samples, highlighting AML-specific and knee-specific loops (**Figure 1B**. We found that approximately 54% of these loops (46,630 of the 86,303 total) are found in both AML and femur samples. Both AML and femur samples showed high heterogeneity between called loops. We found that AML samples had more common loops than femur samples. 5% (1,450 of 29,844 total) of loops in AML samples are common between all three. In comparison, a more limited 2% (183 of 9,829 total) of loops in femur samples are common between all three. Genes near these subsets of loops were also identified (**Table S4**). From the genes associated with loops common to all 3 AML samples, 29 prominent oncogenes including *MYC*, *ABL1*, *RARA*, and *RAD21* were identified. Protein class over-representation analysis of the same gene list revealed over 2-fold enrichment of RNA processing and splicing (*P* = 0.0013 and 0.0026, respectively) (**Figure S2A**). Reactome pathway over-representation analysis of this gene list additionally revealed strong enrichment of various gene sets, including genes involved in cohesin loading onto chromatin (*P* = 0.0029) **(Figure S2B**). Taken together, AML samples show increased loops near key oncogenes, as compared to normal hematopoietic stem cells.

**Figure 1.**
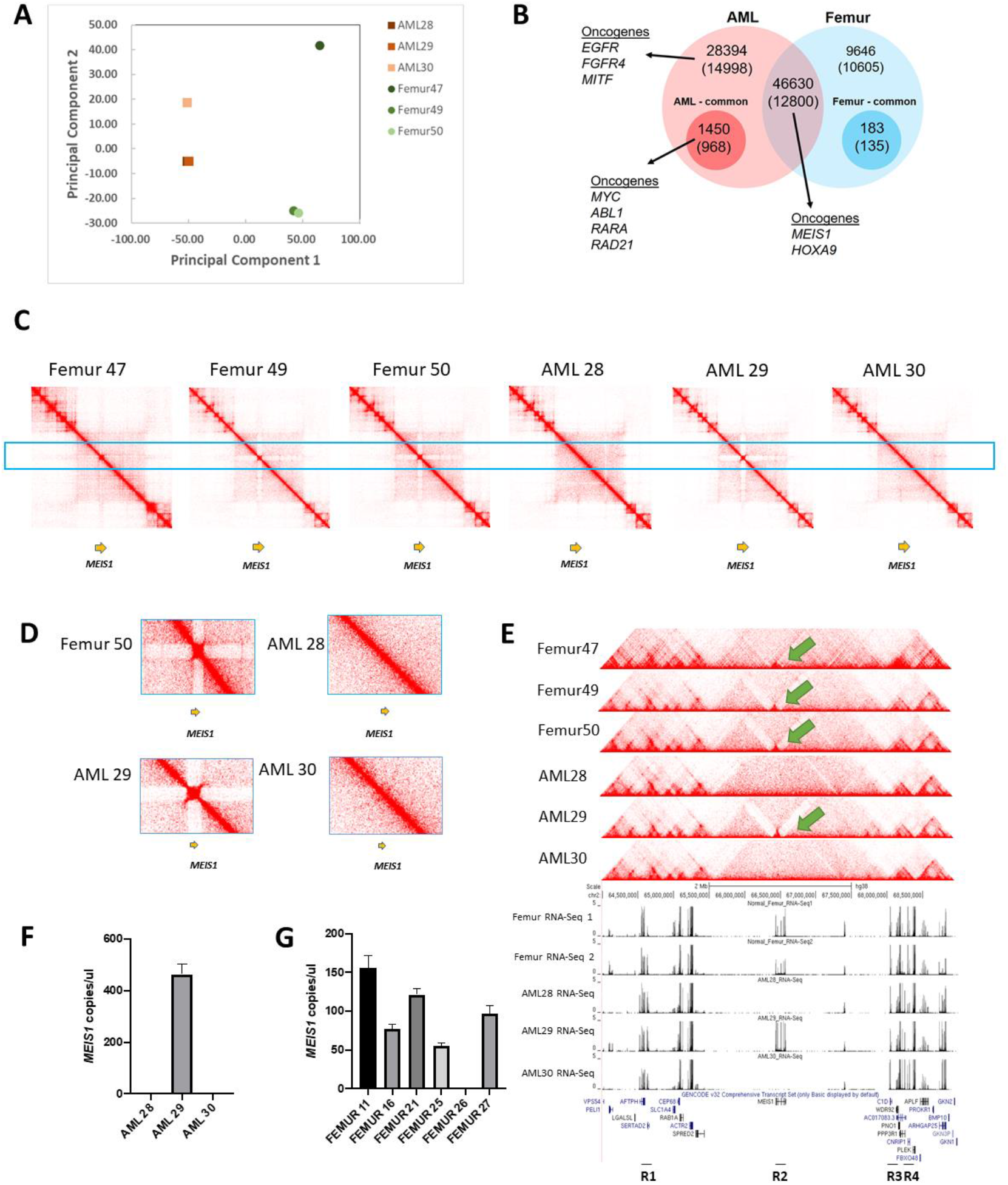
Analyses of chromatin interactions at the *MEIS1* region. **A.** Principal Component Analysis (PCA) plot of AML and normal CD34+ femur samples. **B.** Representation of oncogene profiles in AML and normal femur clinical samples. **C.** Hi-C heatmaps of femur 47, 49, 50 & AML28, 29 and 30 indicate that AML 28 & 30 has an unusual loss of a sub-TAD (genomic coordinates - chr2:64000000-69000000). **D.** Zoomed in view of FIRE region of the clinical samples **E.** Screenshot on the UCSC genome browser of the sub-TAD in alignment to their expression levels at the *MEIS1* gene (genomic coordinates - chr2:64000000-69000000). Coverage normalization was used to visualize the Hi-C heatmaps. **F.** ddPCR of *MEIS1* in CD34+ cells of clinical AML samples. **G.** ddPCR of *MEIS1* in CD34+ cells of clinical normal bone marrow samples.

Hi-C analyses are capable of revealing translocations (22), and therefore we analysed the Hi-C data for structural variants in these patient samples by Dovetail Genomics’ “Selva” approach for analysing structural variants (**Supplementary Methods**). AML28 and AML29 showed fewer than 5 translocations and no known oncofusions from Hi-C analyses, while AML30 showed a common t(8; 21) translocation which fuses *AML1* (also called *RUNX1*) with the *ETO* gene (also called *RUNX1T1*) (**Table S5**).

#### Identification of a Frequently Interacting Region (FIRE) at MEIS1 in haematopoietic stem cells and specific AML samples

Next, we examined chromatin interactions at a key AML oncogene, *MEIS1*. Analysis of the Hi-C heatmaps indicated that all the femurs and AML29 have an unusual region consisting of many interactions in a local space at *MEIS1* (small, tight, dense square shown in the Hi-C heatmap in **Figure 1C-D**). This is absent in AML28 and AML30 (**Figure 1C-D**). The interacting structure encapsulates the *MEIS1* gene from promoter to terminator. We provided a guide to interpret Hi-C heatmaps in **Figure S4**.

The gene expression levels of *MEIS1* in AML28, AML30 and AD796 as examined by RNA-Seq (23) (24) are also observed to be lower than those of normal CD34+ haematopoietic stem cells, while the gene expression level of *MEIS1* in AML29 is much higher than those of normal CD34+ haematopoietic stem cells (**Figure 1E-G & 2C**). Gene expression analysis by digital droplet polymerase chain reaction (ddPCR) of *MEIS1* shows that AML28 and AML30, which have both lost the tight loop structure region, have lower levels of *MEIS1* as compared with AML29 (**Figure 1E**). ddPCR analysis in normal CD34+ haematopoietic stem cells from six femur samples showed that *MEIS1* expression is present in five of six patients, but at a lower level as compared with AML29 (**Figure 1F**).

**Figure 2.**
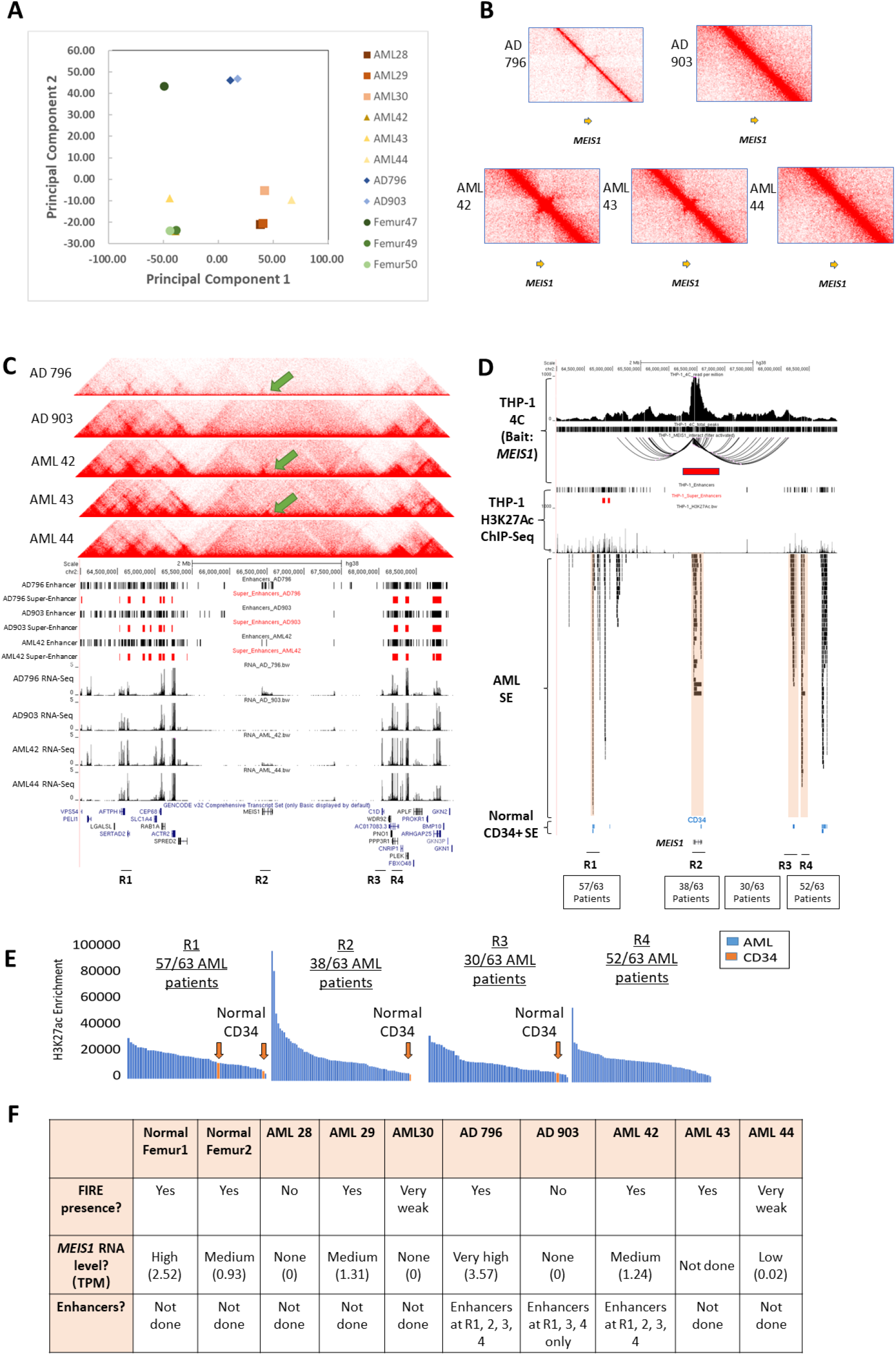
Super-enhancers and the associated chromatin interactions at *MEIS1* FIRE. **A.** Principal Component Analysis (PCA) plot of total AML samples and three normal femur CD+ cells samples. **B.** Hi-C heatmaps of additional clinical AMLCD34+ samples at the *MEIS1* region. (Genomic coordinates: chr2:64000000-69000000) **C.** Aligned Hi-C heatmaps with super-enhancer and gene expression profiles of additional clinical AML CD34+ samples at the *MEIS1* FIRE. **D.** Aligned 4C and H3K27 acetylation signal profile of the THP-1 cell line with the super-enhancer profiles of 63 clinical AML samples. The red box represents the location of the *MEIS1* FIRE. **E.** Ranking of H3K27 acetylation signals for 63 AML clinical patients’ samples and 2 normal CD34+ clinical samples at four super-enhancer regions. **F.** Overview of the FIRE, *MEIS1* expression and enhancer profiles in the clinical samples.

#### Integrated Hi-C, RNA-Seq and H3K27ac ChIP-Seq analysis in AML total bone marrow samples shows chromatin loops that connect distant super-enhancers to the MEIS1 gene promoter

To investigate the heterogeneity of the chromatin interactions at *MEIS1* in more samples, we performed Hi-C on five additional total bone marrow isolates from AML patients (“AML 42”, “AML 43”, “AML 44”, “AD796”, “AD903”). AD796 and AD903 were obtained from total bone marrow that had been previously frozen and were thawed for the experiment, while AML42, 43 and 44 came from fresh isolates. Analysis of the Hi-C heatmaps indicated that AML 42, AML 43 and AD796 have the chromatin interaction structure (**Figure 2B-C, Figure S3A-B**), and this is absent in AML44 and AD903 (**Figure 2B-C, Figure S3A-B**). The quality of the Hi-C library is not as high and the maps are not as distinct for AD796, which may be because the sample was previously frozen and then thawed. Nevertheless, the chromatin interaction structure still can be seen in this sample (**Figure 2B**).

We successfully obtained RNA-Seq libraries in AD796, AD903, AML42 and AML44. AD796 and AML42, which have the chromatin interaction structure, show expression of *MEIS1* (**Figure 2C**). By contrast, AML903 and AML44, which do not have the chromatin interaction structure, do not show expression of *MEIS1* (**Figure 2C**), which demonstrates that the presence of the chromatin interaction structure in *MEIS1* is associated with gene expression of *MEIS1*.

Given that the small TAD-like structure appears in the context of a larger TAD structure but possess chromatin interactions that are unconfined within the small TAD-like structure, we reason that this pattern is characteristic of a recently-reported novel class of chromatin interactions called “Frequently Interacting Regions” (“FIRES”), which are regions of unusually high local contact frequency that tend to be depleted near TAD boundaries and enriched towards the center of TADs (18). Examination of the heatmap in **Figure 1E** and **2C** reveals that the FIRE is located away from the boundary and towards the center of a large TAD. Moreover, FIRES are often reported to be located near cell identity genes and *MEIS1* is important in maintaining stem cell-like identities (25), supporting the notion that the region of interest is likely to be a FIRE. Further details about FIREs are shown in **Figure S4A-B**.

One phenomenon of FIREs is their tendency to be enriched for super-enhancers (SEs). To investigate enhancers and SEs, we performed H3K27ac ChIP-Seq on AD796, AD903 and AML42, and called typical enhancers as well as SEs from the H3K27ac ChIP-Seq data. Interestingly, we observed enhancer presence near the *MEIS1* locus in the AML samples AD796 and AML42, which have the putative *MEIS1* FIRE (**Figure 2C**). By contrast, AD903, which lacks the *MEIS1* FIRE, does not have enhancers near the *MEIS1* locus (**Figure 2C**). This is in line with the RNA-Seq data showing that there is *MEIS1* gene expression in AML796 and AML42, but not AD903 (**Figure 2C**).

Notably, all three samples had SEs within a few megabases of the *MEIS1* region (**Figure 2C**), indicating that the presence or absence of these upstream and downstream SEs is not dependent on the *MEIS1* FIRE. In AML42, which was a sample containing the *MEIS1* FIRE for which we have integrated Hi-C, RNA-Seq and H3K27ac ChIP-Seq data, we could observe Hi-C contacts in the Juicebox matrix that connect upstream and downstream SEs to the *MEIS1* FIRE by chromatin interactions (**Figure 2C, Figure S5A**). These results indicate that the SEs can loop to *MEIS1* via chromatin interactions in an AML sample that contains the *MEIS1* FIRE. By contrast, AD903, which is a sample which lacks the *MEIS1* FIRE and for which we have integrated Hi-C, RNA-Seq and H3K27ac ChIP-Seq data, does not show any loops connecting the *MEIS1* promoter and the distant SEs (**Figure S5B**). Taken together, we speculate that presence of *MEIS1* FIRE and FIRE-associated loops may contribute to *MEIS1* expression via chromatin interactions between the SEs and the *MEIS1* promoter (**Figure S5C**).

Next, we wished to characterize chromatin interactions in the *MEIS1* region and upstream and downstream region in detail. We turned to the THP-1 cell line for this analysis because it possesses a FIRE at the *MEIS1* region (**Figure S6A**). Using Circular Chromosome Conformation Capture (4C) (26,27) in the THP-1 AML cell line with the *MEIS1* promoter as a bait, we found that many interactions to *MEIS1* promoter occur within the FIRE region, confirming that the region indeed consists of many interactions. Moreover, while most interactions occur within the structural region, there are a few chromatin interactions that extend outwards to additional regions within the larger TAD structure (**Figure 2D**). These are the same locations that have super-enhancers in **Figure 2C**. Additionally, 4C analysis of the *MEIS1* promoter in HL-60, another AML cell line, also shows similar chromatin interactions as those found in the THP-1 AML cell line (**Figure S7**).

We investigated how common are super-enhancers around the *MEIS1* locus in AML samples using a dataset of H3K27ac ChIP-Seq libraries that had been performed on total bone marrow from 66 AML patients and 2 healthy individuals (19) to identify AML-acquired SEs. We found that approximately 87% (51 out of 58) of AML patient show the presence of strong H3K27ac ChIP-Seq signals at the *MEIS1* FIRE region **(Figure S6A)**, suggesting the presence of active transcriptional machinery within the FIRE. Moreover, we found four SEs (SE 1, 2, 3 & 4) that are located at the *MEIS1* regions R1, R2, R3 and R4. SE1, SE3 and SE4 are located upstream and downstream of the *MEIS1* gene and can be found in about 79% (57 out of 63) and approximately 47% (30 out of 63) of AML patient samples. SE2, which is located within the *MEIS1* intron, can be found in approximately 60% (38 out of 63) of the AML patients. SE1 was found in two different clinical samples of healthy CD34+ haematopoietic stem cells, and SE2 and SE3 were both present in one of two clinical samples (**Figure 2C-D**), suggesting that several SEs that drive *MEIS1* at high levels are present in haematopoietic stem cells and retained in certain AML patients.

Moreover, the majority of AML samples that had super-enhancers at regions R1, R2, R3 and R4 showed H3K27ac signals at these regions that were higher than the healthy blood stem cells (**Figure 2E & Figure S8**), indicating that SEs originally present in haematopoietic stem cells may further increase in strength upon cancer formation and progression. These extremely strong SEs present in AMLs may explain why the level of *MEIS1* expression is much higher in AML samples with the *MEIS1* FIRE as compared with normal blood stem cells, which also contain the *MEIS1* FIRE.

Taken together, our results indicate that four of eight AML samples examined had the FIRE in *MEIS1*. We obtained RNA-Seq data from three of these four samples, and these three samples showed *MEIS1* gene expression, while the other four AML samples (which did not have the *MEIS1* FIRE) showed absent *MEIS1* gene expression. The *MEIS1* FIRE was associated with enhancers at region 2, but not associated with whether there were super-enhancers present at regions 1, 3 and 4. However, in the Hi-C analysis of a sample with the *MEIS1* FIRE (AML42), chromatin contacts could be seen that connected the super-enhancers to the *MEIS1* promoter, while in the Hi-C analysis of a sample without the *MEIS1* FIRE (AD903) no such contacts were seen (**Figure 2F**).

Next, we asked whether the *MEIS1* FIRE could be found in other cell types. Juicebox (21,28) images of the region in GM12878 human lymphoblastoid cell line show that this structure does not exist in GM12878 lymphoblastoid cells. Additionally, Juicebox images from Hi-C data from (29) of the region around the *MEIS1* gene in mice also shows a FIRE, which is present in mouse embryonic stem cells, lost in differentiated B cells, and re-gained in activated B cells (**Figure S9A**). Moreover, Juicebox images from Hi-C data of the region in T-Acute Lymphocytic Leukemia (T-ALL) show the FIRE in one of four samples (30) (**Figure S9B**).

We observed that this *MEIS1* FIRE structure exists in the cell lines of different tissue types such as K562 (Chronic Myelogenous Leukemia cell line), HUVEC (human umbilical vein endothelial cells), HMEC (human mammary epithelial cells), IMR90 (human fetal lung fibroblast cell line), HeLa (human cervical carcinoma cell line), HAP1 (human near-haploid Chronic Myelogenous Leukemia cell line) and THP-1 (AML cell line) (**Figure S6A**). Gene expression analysis from the Human Protein Atlas showed that most of these cell lines express *MEIS1* either highly (HeLa), moderately (HAP1, HUVEC, THP-1 and K562) or expression unknown in case of HMEC and IMR90 (Figure S6B)(31). Interestingly, we found that HUVEC which moderately expresses *MEIS1* shows strong H3K27ac as well as H3K27me3 ChIP-Seq signals in the FIRE region, raising the possibility that *MEIS1* FIRE could potentially be associated with bivalent chromatin marks in these cells (**Figure S10**) (32) and might explain the moderate levels of *MEIS1* in HUVEC. Other cell types such as Human Mammary Epithelial Cells (HMEC) do not show strong H3K27me3 ChIP-Seq signals, but do not show strong H3K27ac ChIP-Seq signals either, indicating that the FIRE may be present but not associated with active transcription (**Figure S10)**. This suggests that while other cell types besides haematopoietic stem cells have the FIRE, the FIRE may be associated with repressive marks and/or lack activating marks, which keeps *MEIS1* silent.

#### Maintenance of a Frequently Interacting Region (FIRE) boundary near MEIS1 is important for high MEIS1 gene expression in the K562 cell line

Next, we investigated the consequences of perturbation of the *MEIS1* FIRE through CRISPR deletion of a FIRE border. FIRE formation is partially dependent on CTCF (18), and we observed that the 3’ end of *MEIS1* which is near a border of the FIRE is marked by a strong CTCF binding site located at a CpG island that can be found in multiple cell lines. We asked whether maintenance of the CTCF binding site was important in maintaining high levels of *MEIS1* gene expression and proceeded to excise the CTCF binding site near the FIRE boundary in K562 cells (**Figure S11**) via CRISPR.

Our 4C results of the CTCF excised K562 cells as compared with empty vector control cells show a clear loss of *MEIS1-*promoter linked chromatin interactions within the *MEIS1* locus. Furthermore, we see an apparent loss of *MEIS1* associated distal chromatin loops that were beyond the immediate CTCF boundaries of *MEIS1* (**Figure 3A**), including losses of chromatin interactions between the *MEIS1* promoter and the SEs (SE1, 2, 3 & 4) **(Figure 3A).** Taken together, our results indicate that excision of the CTCF binding site at the border of the *MEIS1* FIRE can lead to severe loss of chromatin interactions.

**Figure 3.**
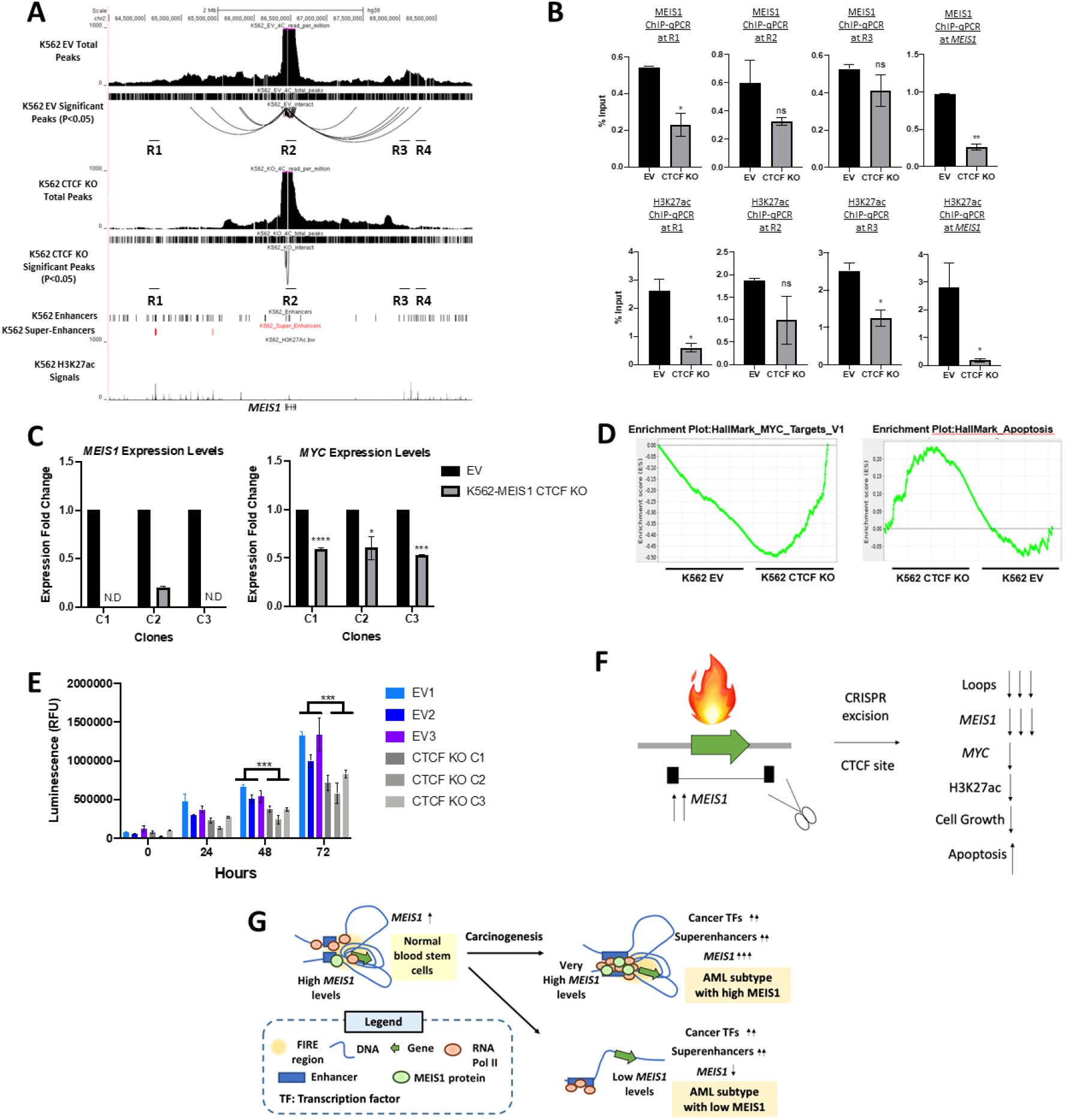
CRISPR excision of a CTCF binding site near the *MEIS1* FIRE leads to loss of *MEIS1.* **A.** 4C of K562 EV (empty vector) and CTCF KO (Knock-out) are shown in comparison with their total peaks and significant loops (P<0.05) (Genomic Coordinates: chr2:64000000-69000000). The locations of the SEs (super-enhancers) are marked as R1-4. **B.** Comparison of MEIS1 and H3K27ac enrichment profiles at R1, R2, R3 and the *MEIS1* promoter in K562 EV (empty vector) and CTCF KO (Knock-out) cells. Data shown are average +/− standard error from two biological replicates. **C.***MEIS1* RT-qPCR (left) and *MYC* RT-qPCR in negative control cells which went through the CRISPR process with EV (“empty vector”) and CRISPR KO (“CRISPR Knockout”) cells. Data shown are average +/− standard error from three biological replicates. “N.D.” indicates no gene expression was detected. One-tailed t-test was performed to evaluate significance, and the asterisks indicate the data is significant as per ****P<0.0001, *** - P< 0.001 ** - P< 0.01, *- P < 0.05 level. D. Gene Set Enrichment Analysis (GSEA) of RNA-seq profiles from K562 empty vector and K562 CTCF KO cells. **E.** Cell viability assays on three clones of K562 EV (empty vector) and three clones of K562 CTCF KO (Knock-out) cells. Data shown are average +/− standard error from three biological replicates. Two-tailed T-test was performed to evaluate significance, and the asterisks indicate the data is significant as per *** - P< 0.001. **E.** Schematic summary of the data presented. CRISPR excision of the FIRE (indicated by the “fire” icon) at *MEIS1* leads to *MEIS1* gene expression and other cellular changes in myeloid leukemia. **F.** A proposed schematic of the potential mode of carcinogenesis in high *MEIS1* (*MEIS1* FIRE present) and low *MEIS1* (*MEIS1* FIRE loss) expressing myeloid leukaemias.

We next performed ChIP-qPCR on the K562 empty vector and CTCF Knockout cells to assess the MEIS1 and H3K27ac enrichment levels at the four regions (R1-4) as well as at the *MEIS1* promoter (**Figure 3B and S12**). We investigated MEIS1 binding to super-enhancers and the MEIS1 promoter because we reasoned that MEIS1, as a transcription factor, might auto-regulate. Our ChIP-qPCR results showed that the MEIS1 indeed binds to super-enhancers and the *MEIS1* promoter in K562 cells (**Figure 3B**). Additionally, MEIS1 enrichment levels are significantly reduced at R1 that is suggested to possess super-enhancers and the *MEIS1* promoter upon the CTCF knockout in the K562 cells. In addition, the H3K27ac enrichment levels were significantly downregulated at R1, R3 and the *MEIS1* promoter (**Figure 3B**). However, the H3K27ac enrichment levels were not significantly different at R2 (**Figure 3B**), and the H3K27ac enrichment levels were significantly upregulated at R4 (**Figure S12**). These results suggest that the loss of the *MEIS1* CTCF region can lead to alterations in MEIS1 and H3K27ac enrichment at the regions that possesses SEs and *MEIS1* promoter regions. Most of the super-enhancers showed reduced MEIS1 binding and H3K27ac enrichment, suggesting a weakening of the enhancer ability of these regions, however, R4 showed an opposite trend, indicating that the landscape of enhancer changes is complex.

Examination of *MEIS1* by reverse transcriptase quantitative polymerase chain reaction (RT-qPCR) in cells treated with empty vector showed high levels of *MEIS1*, but a marked reduction of *MEIS1* was observed in the CTCF CRISPR knockout cells (**Figure 3C**). *MYC*, which was previously shown to be a downstream target of *MEIS1* in zebrafish (33), also showed significantly lower gene expression in CTCF CRISPR knockout cells (**Figure 3C**). The CTCF CRISPR knockout cells also grew more slowly in culture as compared to the control empty vector K562 cells (**Figure 3E)**. Interestingly, Gene Set Enrichment Analyses (GSEA) performed on the RNA-Seq analyses of the K562 EV and CTCF KO cells also showed a reduction in the expression of MYC target genes and increased expression of apoptosis-related genes in K562 cells upon CTCF knockout at the *MEIS1* FIRE. These suggest that the reduction in MYC expression from the loss of *MEIS1* may result in the reduced expression of MYC target genes (Figure 3D). Overall, we found that maintenance of the CTCF sites at the FIRE is important in maintaining H3K27ac enhancer levels, *MEIS1* gene expression levels and cell growth in myeloid leukemia cells (**Figure 3E**).

Taken together, our results suggest a model of *MEIS1* gene regulation in which blood stem cells have a FIRE at *MEIS1* that shows many chromatin interactions to other genomic regions, which can be occupied by SEs. The interplay between SEs and the FIRE may drive *MEIS1* at high levels in blood stem cells and AML samples with intact FIREs. In AMLs that lack the FIRE and chromatin interactions between the FIRE and the SEs, we speculate that the absence of contacts between the MEIS1 promoter and super-enhancers leads to lower *MEIS1* expression. These AMLs that have low *MEIS1* levels may be driven by other AML oncogenes, and thus do not rely on *MEIS1* gene expression for survival. Therefore, the presence of different chromatin conformations in various AMLs at *MEIS1* and their varying interactions with SEs may explain the heterogeneity of *MEIS1* expression (**Figure 3F-G**).

#### Genomic loci that can loop to the HOXA9 oncogene display heterogeneity in SE occupancy in AMLs

Next, we investigated *HOXA9*, an oncogenic transcription factor that tends to be co-expressed with *MEIS1*. *HOXA9* is expressed at a moderately high level in CD34+ normal haematopoietic stem cells and overexpressed in half of AMLs (**Figure 4A)**. To characterize the heterogeneity of *HOXA9* levels in AML, we performed digital droplet PCR to compare *HOXA9* levels with progenitor (CD34+ haematopoietic stem cells) and more differentiated myeloid cells (CD33+ myeloid cells) derived from healthy bone marrow samples. We found *HOXA9* expression levels to be higher in CD34+ cells, while lower *HOXA9* expression levels were observed in myeloid cells. By contrast, AML samples exhibited a range of *HOXA9* expression levels; from very low levels to very high levels but all of these expression levels are higher than those of the CD34+ cells from femurs (**Figure 4A**). Next, we performed ddPCR of *HOXA9* on AML28, AML29 and AML30 (**Figure 4B**), as well as six other CD34+ enriched cell populations from healthy patients (“Femur11”, “Femur16”, “Femur21”, “Femur25”, “Femur26”, “Femur27”) (**Figure 4C**). We found that all the normal cells had varying levels of *HOXA9*, while AML29 had high *HOXA9* gene expression and AML28 and AML30 had low *HOXA9* gene expression (**Figure 4B & 4C**).

**Figure 4.**
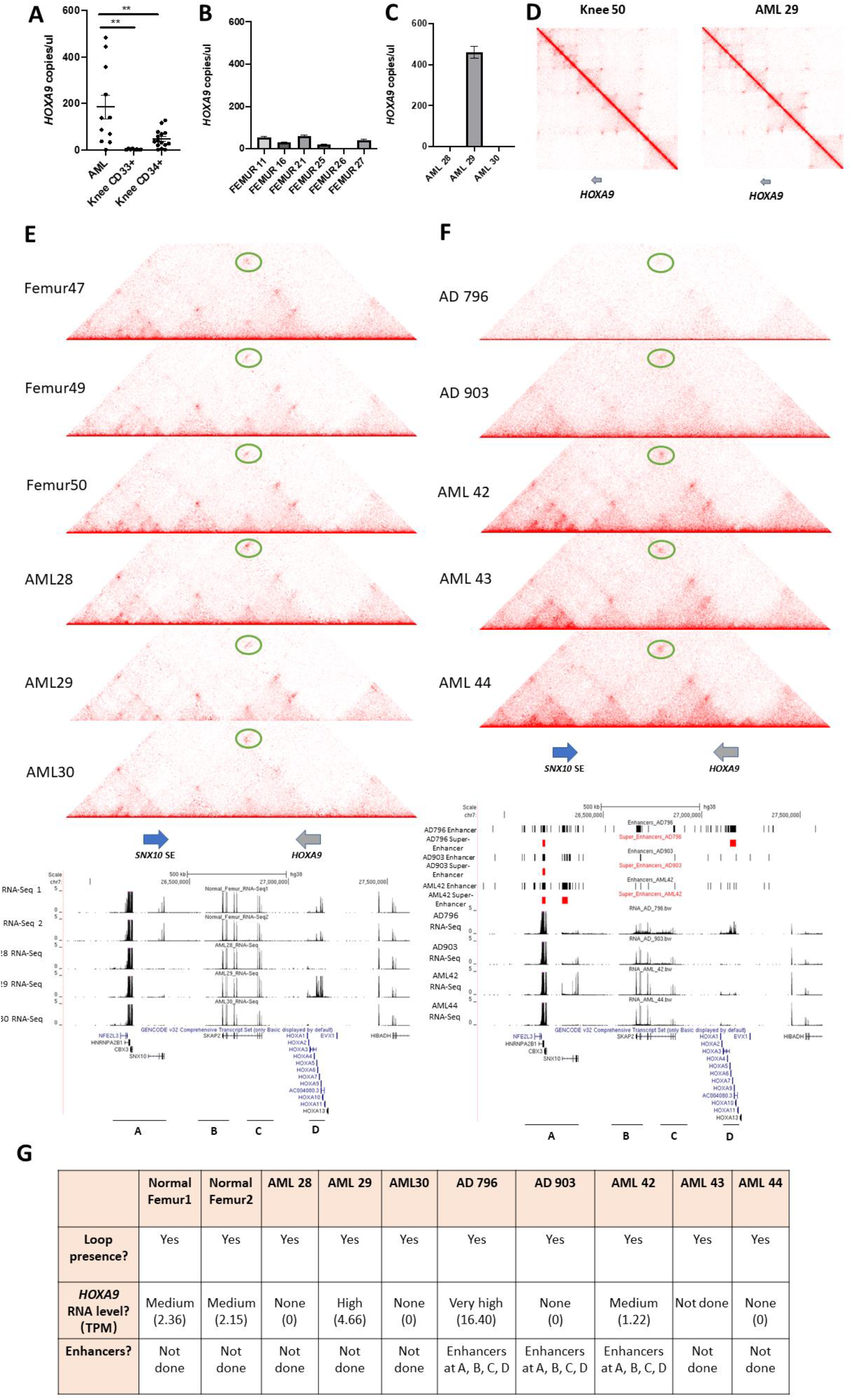
Hi-C analyses of CD34+ blood stem cells from Acute Myeloid Leukemia patient samples and normal femur bone marrow shows abundant loops and Topologically Associated Domains (TADs) around the *HOXA9* gene. **A.** Digital droplet (ddPCR) investigations of *HOXA9* gene expression level in AML, CD33+ myeloid cells from healthy donors, and CD34+ cells from healthy donors. Significance testing was performed by Student’s two-tailed t-test. Two asterisks ** indicate p<0.01. **B.** ddPCR of *HOXA9* in CD34+ femur 25, 26 and 27 samples. Data shown indicates the average value of technical replicates performed on the same clinical sample and error bars indicate standard error. **C.** ddPCR of *HOXA9* in CD34+ AML28, 29, 30 samples. Data shown indicates the average value of technical replicates performed on the same clinical sample and error bars indicate standard error.**D.** Heatmaps (coverage normalized) of the *HOXA9* genomic region in Femur 50 (left) and AML29 (right) depicted have the X axis that list a set of genomic coordinates (in the case of the *HOXA9* region it is: chr7:25866917-27612715). The same coordinates are then shown on the Y axis, and regions with strong interactions are shown in dark red. Loops are indicated as dots on the map. Distinct squares indicating regions with strong interactions can be seen in the heatmap, indicating TADs which are regions of the genome with strong interactions within the TAD boundaries. **E.** The heatmap shows the TADs of three normal femur bone marrows and three AML derived CD34+ samples. The loop indicated in the heatmap is depicted (as a dot), and the loop depicted is marked by a red circle in the screenshot. The genomic locations of the Region A (*SNX10)* and *HOXA9* genes in this area are indicated. The RNA-seq tracks show the expression levels of *HOXA9* and the *SNX10* SE in two normal CD34+ femur samples and the three AML CD34+ samples. **F.** The heatmap shows the TADs of five AML derived CD34+ samples. The loop indicated in the heatmap is depicted (as a dot), and the loop depicted is marked by a red circle in the screenshot. The genomic locations of the Region A (*SNX10*) and *HOXA9* genes in this area are indicated. The RNA-seq tracks show the expression levels of *HOXA9* and the *SNX10* SE in four AML CD34+ samples (AD796, AD903, AML 42 & AML44). Super-enhancer and enhancer presence in three AML CD34+ samples (AD796, AD903 & AML 42) at the *HOXA9* locus and Region A (*SNX10* SE) are also aligned to their Hi-C heatmaps and RNA-seq profiles.

Looking at Juicebox images around *HOXA9* from our Hi-C data, we found strong chromatin interactions and/or TADs around the *HOXA9* gene (**Figure 4D & Figure S13A & S13B**). In contrast to *MEIS1*, the *HOXA9*-associated chromatin interactions and TADs were found in all femur samples as well as AML samples (**Figure 4E & 4F**), indicating that different chromatin interaction usage was not the reason behind the different levels of *HOXA9* oncogene expression in these cells.

We hypothesized that differences in superenhancer involvement with chromatin interactions might underlie the different levels of *HOXA9* expression instead. RNA-Seq of the clinical AML and normal CD34+ samples show that *HOXA9* expression was higher in both normal femurs derived CD34+ cells and in three AML samples (AML 29, AD796 & AML 42), of which two of these AML samples possess enhancers in the regions with chromatin interactions to *HOXA9*. Based on our integrated analyses of Hi-C and H3K27ac ChIP-seq analysis, we observed the presence of H3K27ac signals at three regions (Region A, B & C) upstream of the *HOXA9* gene (Region D). These three regions also form chromatin interactions to the *HOXA9* gene locus, thus suggesting that super-enhancers/enhancers may be present at this region to regulate the *HOXA* gene cluster in AML. Interestingly, we observed that *HOXA9* expression level is distinctly higher in the AML sample, AD796 which possess strong levels of enhancers and super-enhancer presence at the *SNX10* SE and the *HOXA9* region, while the lack of strong enhancers at *SNX10* SE and *HOXA9* region resulted in lower *HOXA9* expression levels (**Figure 4F**).

We then investigated AML cell lines that have high or low levels of *HOXA9* oncogene expression. ddPCR analysis of *HOXA9* expression levels in THP-1 and HL-60 AML cell lines indicate that *HOXA9* is specifically upregulated in THP-1 but not HL-60 (**Figure S14**). We investigated the chromatin interactions using 4C with the *HOXA9* gene promoter as the bait region in THP-1 and HL-60 myeloid leukemic cell lines (**Table S8 & Figure S15**). We found that the chromatin interactions were similar between HL-60 and THP-1 cell lines (**Figure 5B**), confirming that different chromatin interaction usage was not the reason behind the different levels of *HOXA9* oncogene expression in these cells.

**Figure 5.**
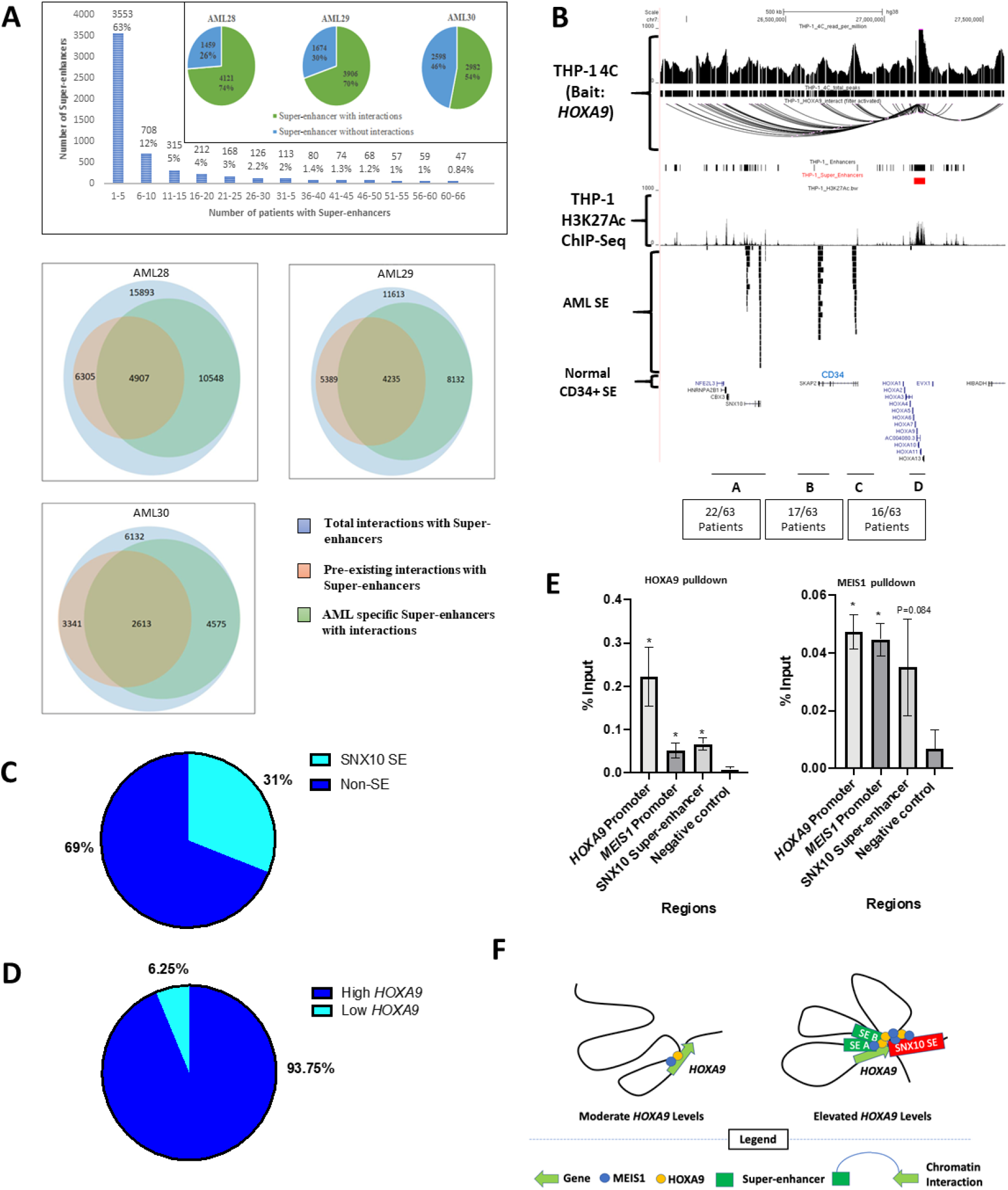
An AML-specific SE hijacks pre-existing chromatin interaction between the *SNX10* gene and *HOXA9* and is associated with high *HOXA9* levels. **A.** Chart depicting the number of SEs found in 66 AML patients. Individual boxes indicate the SE profiles of three AML samples. Pie chart shows the number of pre-existing SE associated chromatin loops in AML. **B.** The 4C loops and H3K27ac profile of THP-1 are shown in alignment with SE profiles from two normal CD34+ donors and 63 AML patients (each row represents one patient) at the location between Region A (*SNX10)* and *HOXA9*. The black bars indicate SEs. Genomic coordinates for region Chr7:25,906,537-27,652,334. **C.** Pie chart showing the proportion of AML patients from the 66 AML patients who have the *SNX10* SE (“SNX10 SE”) or not (“no SE”). **D.** Pie chart showing the proportion of AML patients with the *SNX10* SE with high *HOXA9* as compared with low *HOXA9*. **E.** Chromatin-immunoprecipitation (ChIP)-qPCR from HOXA9 and MEIS1 antibody pulldown at selected oncogenes promoter regions. **F.** Schematic of the hypothesized mechanism of SEs ( at Regions A, B & C) hijacking pre-existing chromatin loops at the *HOXA9* locus.

To explore the hypothesis that SE acquisition in AML might be heterogenous, with some patients acquiring certain SEs and not others, possibly leading to the heterogeneous overexpression of *HOXA9* seen in some AML patients, we examined the patient SEs identified earlier from McKeown *et al.* (19). We found that 63% of SEs could be found in fewer than 5 patients, and fewer than 2% of them could be found in more than 55 of 66 patients, indicating that SE acquisition in AML is heterogeneous (**Figure 5A**).

To characterize the interplay between SEs and chromatin interactions, we intersected the SE information with the chromatin loops detected in the normal and AML Hi-C data. In AML28, we observed that 74% of SEs were associated with chromatin interactions (**Figure S16A**). In AML29, 70% SEs were associated with chromatin loops. In AML30, 54% SEs were associated with chromatin loops (**Figure 5A**). Some of the observed SEs connect to genes with potential roles in cancer via chromatin interactions. For example, an AML SE can connect with *PKP4* gene, known to be differentially expressed in human lung squamous cell carcinoma (34) through two chromatin interactions (**Figure S16B**).

We further observed that approximately half of the chromatin interactions associated with AML-specific SEs could be also found in normal CD34+ haematopoietic stem cells (4907 pre-existing interactions out of 10548 = 46.5% in AML28, 4235 pre-existing interactions out of 8132 = 52.07 % in AML29 and 2613 pre-existing interactions out of 4575 = 57.11% in AML30) (**Table S7)**. This indicates that while some SEs that are newly acquired in certain AML patients might be associated with newly formed chromatin interactions, other SEs that are newly acquired in AML might occupy pre-existing chromatin interactions that are also found in the precursor cells (haematopoietic stem cells) to regulate target genes.

We then investigated SE occupancy of chromatin interactions at *HOXA9*. We detected the presence of multiple SEs (**Figure 5B**) that show interactions with the *HOXA9* promoter through 4C & Hi-C. In particular, we observed a SE at region A (*SNX10* SE) that loops over to *HOXA9* (Region D) (**Figure 5B**), which we analysed further. This *SNX10* SE that is found at region A is specifically present in 18 of 58 of the AML patients and is not found in the normal CD34+ haematopoietic stem cell data from McKeown *et al.* (19). Our analysis revealed that about 31% of AML patients carry this *SNX10* SE (**Figure 5C**). The presence of the *SNX10* SE was associated with upregulated high *HOXA9* levels in the AML patient samples (**Figure 5D**).

Next, because HOXA9 and MEIS1 are transcription factors, we investigated how HOXA9 and MEIS1 bind to SE and promoters in the *HOXA9* and *MEIS1* gene regions. ChIP-qPCR of THP-1 at regions of the *SNX10* SE and the *HOXA9* promoter showed the binding of HOXA9 and MEIS1 proteins at both locations (**Figure 5E)**. Interestingly, we also observed that the HOXA9 and MEIS1 proteins bind to the *MEIS1* promoter regions, suggesting a tight circuit between HOXA9 and MEIS1. Altogether, our results suggest a model whereby the *MEIS1* FIRE can lead to high *MEIS1* levels. The MEIS1 transcription factor can auto-regulate the *MEIS1* promoter, and also bind to the *HOXA9* promoter. HOXA9 can bind to the *HOXA9* and *MEIS1* promoters, as well as the *SNX10* SE that loops over to the *HOXA9* promoter. This can explain the observation that *MEIS1* and *HOXA9* tend to be upregulated together in AML. Additionally, the heterogeneity of super-enhancer acquisition at chromatin interactions that loop to *HOXA9* in AML patients may explain the heterogeneity of *HOXA9* oncogene expression observed in AML patients (**Figure 5F and Figure S17**).

## Discussion

Previous investigations into 3D genome organization in cancers by genome-wide approaches has revealed that there are chromatin interactions at oncogenes in primary gastric adenocarcinoma (35) and acute lymphocytic leukemia (30). However, there have been no Hi-C analyses that have compared chromatin interactions between normal precursor cells and cancer cells. Here we performed Hi-C comparing three samples of CD34+ haematopoietic stem cells from healthy individuals as well as three samples of CD34+ cells from acute myeloid leukemia patients and a further five samples of total bone marrow from acute myeloid leukemia patients. We found that there were more common chromatin interactions to oncogenes in acute myeloid leukemia as compared with healthy individuals, suggesting that chromatin interactions might be a potential therapeutic vulnerability in acute myeloid leukemia.

Additionally, a major focus in understanding cancer is identifying the molecular underpinnings of the heterogeneity of oncogene expression in different cancer clinical samples. We identified two mechanisms involving chromatin interactions that may give rise to heterogeneity of oncogene expression. The first mechanism is that different AMLs may show different patterns of chromatin interactions around oncogenes, thus leading to different oncogene expression levels, such as *MEIS1*, which is regulated by a FIRE. A second mechanism is that SEs may be heterogeneously acquired in AML. The AMLs which have acquired such SEs may show high levels of oncogene expression because the SEs may occupy, or “hijack”, pre-existing enhancer-promoter chromatin interaction circuits present in precursor cells such as haematopoietic stem cells, thus driving oncogenes strongly in certain AMLs but not others.

HOXA9 and MEIS1 have been widely reported to be important contributors to the progression of AML (9,10). The synergistic effect of HOXA9 and MEIS1 also leads to the development of aggressive AML with poor prognosis. Despite the progress in AML research, the understanding of *HOXA9* and *MEIS1* regulation at the epigenomic level remains limited. Here, we discovered auto-regulatory and cross-regulatory chromatin interaction circuits necessary for maintaining high levels of *MEIS1* and *HOXA9* which are seen in some subtypes of leukemia.

Our results demonstrate for the first time that a FIRE at the *MEIS1* region is critical in maintaining high *MEIS1* levels and cancer cell survival. Interestingly, the FIRE at *MEIS1* is not observed in AML28, AML30, AML 44 and AD903 but seen in AML29, AML 42, AML 43 and AD796, suggesting that certain AMLs can develop or be maintained without having FIREs around oncogenes such as *MEIS1*. These AMLs are likely to be driven by different oncogenes or other genetic and epigenetic alterations. For example, AML30 contains a translocation to the *RUNX1* oncogene, suggesting that AML30 does not require high *MEIS1* and *HOXA9* levels to function but might be addicted to *RUNX1* instead.

One question is what could lead to differences in the FIRE presence in different AML samples? One possibility is that there might be genetic differences in the AML samples at the FIRE region. Another possibility is that there might be differences in DNA methylation at the CTCF binding site at the FIRE region. Previous papers have shown that differences in DNA methylation can lead to differential binding of CTCF (36). Additionally, differences in transcription factor binding may lead to differences in FIRE maintenance. In future work, genetic, bisulphite and ChIP-Seq analyses could help to answer this question.

Given that chromatin interactions might be a potential therapeutic vulnerability in Acute Myeloid Leukemia, to further investigate whether targeting a chromatin interaction boundary marked by CTCF, which might lead to changes in chromatin interactions, would lead to reduced oncogene expression, we performed CRISPR excision to perturb a CTCF region at a *MEIS1* FIRE boundary. We showed that this procedure led to altered chromatin interactions and gene expression of *MEIS1*.

One question arising is how can we perturb chromatin interactions in cancer? Aza-cytidine, a drug commonly used in AML treatment, has recently been shown to alter CTCF levels (37). We speculate that these drugs may work in part by leading to alterations in chromatin interactions, such as the FIRE at *MEIS1*, leading to downregulation of gene expression. Moreover, a study characterizing chromatin interactions in Acute Lymphocytic Leukemia patient samples by Hi-C indicates that inhibition of super-enhancers by drugs such as THZ1 can lead to reductions in chromatin interactions at target oncogenes and reduced gene expression (30). Additionally, SE analyses have pinpointed specific oncogenes to which certain AMLs may be addicted, such as retinoic acid receptor alpha (*RARA*), allowing for the development of RARA inhibitors for this category of AMLs (19). Interestingly, RARA was also identified to be associated with common AML-specific loops (Figure 1A) in our Hi-C analysis.

Taken together, our Hi-C maps in this manuscript provide a framework for future investigations of chromatin interactions that are affected by various AML drugs and identify distal oncogenes that are the targets of distal SE that loop over to the gene promoters via chromatin interactions.

In conclusion, we demonstrated that chromatin interactions are altered in AML cells as compared with normal precursor cells. Different chromatin conformations and/or different occupancy by SEs are associated with different levels of oncogene expression. Removal of a FIRE boundary led to reduction of oncogene expression and cancer cell growth. Thus, our results present a molecular basis for the future development of epigenetic inhibitors to target chromatin interactions as a potential therapeutic vulnerability in AML.

## Methods

Detailed methods are given in the **Supplementary Materials**. Briefly, bone marrow samples from AML patients were taken from the back of the pelvic (hip) bone while bone marrow from healthy counterparts was withdrawn during Total Knee Arthroplasty as part of a standard operative procedure. All clinical samples were obtained from the National University Hospital Singapore and collected according to the requirements of the Human Biomedical Research Act. Informed consent was obtained for all clinical samples used in the study. Isolation of CD34+ cells from normal sample mononuclear cells (MNCs) was performed according to the manufacturer’s instructions using CD34 MicroBead Kit UltraPure. Acute Myeloid Leukemia cell lines THP-1 and HL-60 and Chronic Myelogenous Leukemia cells K562 were cultured at 5% CO_2_ at 37^0^C. THP-1 and K562 were cultured with Roswell Park Memorial Institute (RPMI) 1640 media (Hyclone) supplemented with 10% heat-inactivated Fetal Bovine Serum (FBS; Hyclone) and 1% penicillin/streptomycin (Hyclone). HL-60 cells were cultured using Iscove’s Modified Dulbecco’s Medium (IMDM; Gibco), supplemented with 20% heat inactivated FBS (Hyclone) and 1% penicillin/streptomycin (Hyclone). Hi-C was performed through Dovetail Genomics for samples Femur 47, Femur 49, Femur 50, AML 28, AML 29 and AML 30 and by using ARIMA genomics Hi-C kit for samples AD 796, AD 903, AML 42, AML 43 and AML 44. All Hi-C libraries were sequenced by Illumina sequencing. Hi-C data were aligned and processed by Juicer (version 1.5). The reference genome was hg38. RNA was extracted using the DNA/RNA Allprep Kit (Qiagen). Reverse transcription of RNA into cDNA was performed using the qScript cDNA Supermix (Quantabio). Quantitative polymerase chain reaction (qPCR) was performed with the GoTaq qPCR Mastermix (Promega) and QuantStudio 5 Real Time PCR (Applied Biosystems). ddPCR experiments on cDNA were performed with the EvaGreen Mastermix (Biorad) and the QX200 Droplet Digital PCR system. 4C-seq was performed as previously described with some modifications as described in the Supplementary Materials (38). CRISPR-Cas9 excision was performed with the All-in-One vector system as described previously (39). Transfection of K562 cells was performed with the Neon Transfection System (ThermoFisher). All primer sequences are reported in **Table S9**.

## Acknowledgements

This research is supported by the National Research Foundation (NRF) Singapore through an NRF Fellowship awarded to M.J.F (NRF-NRFF2012-054) and NTU start-up funds awarded to M.J.F. This research is supported by the RNA Biology Center at the Cancer Science Institute of Singapore, NUS, as part of funding under the Singapore Ministry of Education Academic Research Fund Tier 3 awarded to Daniel Tenen (MOE2014-T3-1-006). This research is supported by the Singapore Ministry of Health’s National Medical Research Council under its Singapore Translational Research (STaR) Investigator Award, and by the National Research Foundation Singapore and the Singapore Ministry of Education under its Research Centres of Excellence initiative that are awarded to D.G.T. This reserch is supported by NIH grants (R35CA197697 and P01HL131477) that are awarded to D.G.T. This research is supported by a National Research Foundation Competitive Research Programme grant awarded to V.T. as lead PI and M.J.F. as co-PI (NRF-CRP17-2017-02). This research is supported by a Ministry of Education Tier II grant awarded to M.J.F. as PI (T2EP30120-0020). This research is supported by the National Research Foundation Singapore and the Singapore Ministry of Education under its Research Centres of Excellence initiative.

## Author contributions

B.W, L.K, D.B and M.J.F conceived of the research. M.J.F, B.W, L.K, D.B, R.C and D.G.T. contributed to the study design. B.W generated the CRISPR clones and performed 4C, ddPCR, qPCR, ChIP-qPCR and growth assays. L.K performed bioinformatics analysis of the clinical Hi-C, RNA-Sequencing data and ChIP-Seq data. D.B prepared the AML and Knee samples for Hi-C by Dovetail genomics, generated the AML total bone marrow Hi-C libraries and performed manual curation of the Hi-C data. D.B and W.F prepared the clinical samples for RNA-Sequencing. DB prepared the H3K27ac ChIP-Seq libraries on AML MNC samples. R.C performed bioinformatics analysis on published AML cases to identify SEs and their looping patterns in AML. B.W.J.H. performed bioinformatics analyses to understand common and unique loops between AML and femur samples. W.F, J.Q.T and Y.G collected femur and bone marrow samples and isolated CD34+ and CD33+ cells. MCC and WW provided normal bone marrow clinical samples. O.A. and H.Y provided the bioinformatics pipeline and CSI web portal for the 4C analyses. C.Y.T and T.B provided the access for the online storage of clinical data. F.F.S, P.C and C.W.J provided AML clinical samples and annotation. B.W, L.K, D.B, R.C, D.G.T. and M.J.F reviewed the data. B.W, L.K, D.B and M.J.F wrote the manuscript. All authors reviewed and approved of the manuscript.

